# Cortical state transitions and stimulus response evolve along stiff and sloppy parameter dimensions, respectively

**DOI:** 10.1101/839365

**Authors:** Adrián Ponce-Alvarez, Gabriela Mochol, Ainhoa Hermoso-Mendizabal, Jaime de la Rocha, Gustavo Deco

## Abstract

Previous research showed that spontaneous neuronal activity presents sloppiness: the collective behavior is strongly determined by a small number of parameter combinations, defined as “stiff” dimensions, while it is insensitive to many others (“sloppy” dimensions). Here, we analyzed neural population activity from the auditory cortex of anesthetized rats while the brain spontaneously transited through different synchronized and desynchronized states and intermittently received sensory inputs. We showed that cortical state transitions were determined by changes in stiff parameters associated with the activity of a core of neurons with low responses to stimuli and high centrality within the observed network. In contrast, stimulus-evoked responses evolved along sloppy dimensions associated with the activity of neurons with low centrality and displaying large ongoing and stimulus-evoked fluctuations without affecting the integrity of the network. Our results shed light on the interplay among stability, flexibility, and responsiveness of neuronal collective dynamics during intrinsic and induced activity.

## Introduction

How biological systems achieve a tradeoff between stability and flexibility is a central question in biology. A candidate explanation for the coexistence of these two features is *sloppiness* (Machta et al., 2013; Transtrum et al., 2015). In general, sloppiness is a property of complex models exhibiting large parameter uncertainty when fit to data, meaning that different combinations of parameters lead to a similar system behavior, while changes in some few critical parameters, called stiff parameters, significantly modifies it. In this way, biological systems can be either robust to large fluctuations of input/environmental signals which effects are embedded in a high-dimensional subspace of insensitive parameters, or, on the contrary, by tuning some few parameters, configured to be highly sensitive and selective to relevant signals.

Recently, it has been shown that the spontaneous activity of neural circuits presents sloppiness both *in vitro* and *in vivo* (Panas et al., 2015), suggesting that collective activity is stabilized by a subset of highly active and stable neurons, while the activity and co-activity of the remaining neurons present larger spontaneous fluctuations without strongly affecting the collective statistics. However, this view is challenged by extensive research showing that the spontaneous cortical activity transits through different synchronized and desynchronized cortical states (Marguet and Harris, 2011; Harris and Thiele, 2011; Luczak et al., 2013; Pachitariu et al., 2015) that represent statistically different collective behaviors (Hahn et al., 2017) with different information processing capabilities (Pachitariu et al., 2015; Engel et al., 2016; Beaman et al., 2017). Moreover, how sensory inputs affect sloppiness is unknown and it is a relevant question to understand how sensory stimuli change the network state in a way that responsiveness and stability are ensured. In the present study, we examined how changes in neural network parameters correlate with spontaneous transitions among cortical states and stimulus-evoked responses.

To answer these questions, we recorded the neuronal spiking activity in the primary auditory cortex (A1) of six anesthetized rats. We analyzed the joint activity of groups of neurons while the cortex spontaneously transited through different synchronized and desynchronized cortical states and intermittently received external acoustic stimuli. We used a statistical model to describe the joint spiking activity with a small number of parameters. We found that the estimated parameters of neuronal ensemble activity presented sloppiness and that sensory inputs and cortical state transitions evolved in different pathways in parameter space. Specifically, we found that cortical state transitions evolve along stiff dimensions, whereas sensory-evoked activity evolves along sloppy dimensions. Finally, we showed that stiff parameters are related to the activity and co-activity of neurons with high centrality within the functional network of the recorded neurons.

## Results

We recorded spontaneous and stimulus-evoked population activity from the primary auditory cortex (A1) of urethane-anesthetized rats (n = 6) using multisite silicon microelectrodes (see **Methods**). The data was composed of activity from well-isolated single units (44-147 neurons) and some spike-trains from multi-unit activity (3-103 spike-trains). Unless otherwise specified, the analyses present here focused on single-unit activity only. We analyzed the data during spontaneous activity and in response to acoustic “clicks” (5-ms square pulses; inter-stimulus interval, 2.5 or 3.5 s). To track the evolution of the neuronal activity, we divided each recording session into *N_E_* adjacent epochs of 100 s, each one containing 12–29 stimulus presentations. Within each 100-s epoch the data was separated into spontaneous activity, i.e., the activity during 1.5-s intervals preceding each stimulus (i.e., 18–43.5 s of spontaneous activity in total for each epoch), and stimulus-evoked activity, i.e., the activity right after the stimulus onset (**Figure 1A–B**).

**Figure 1.**
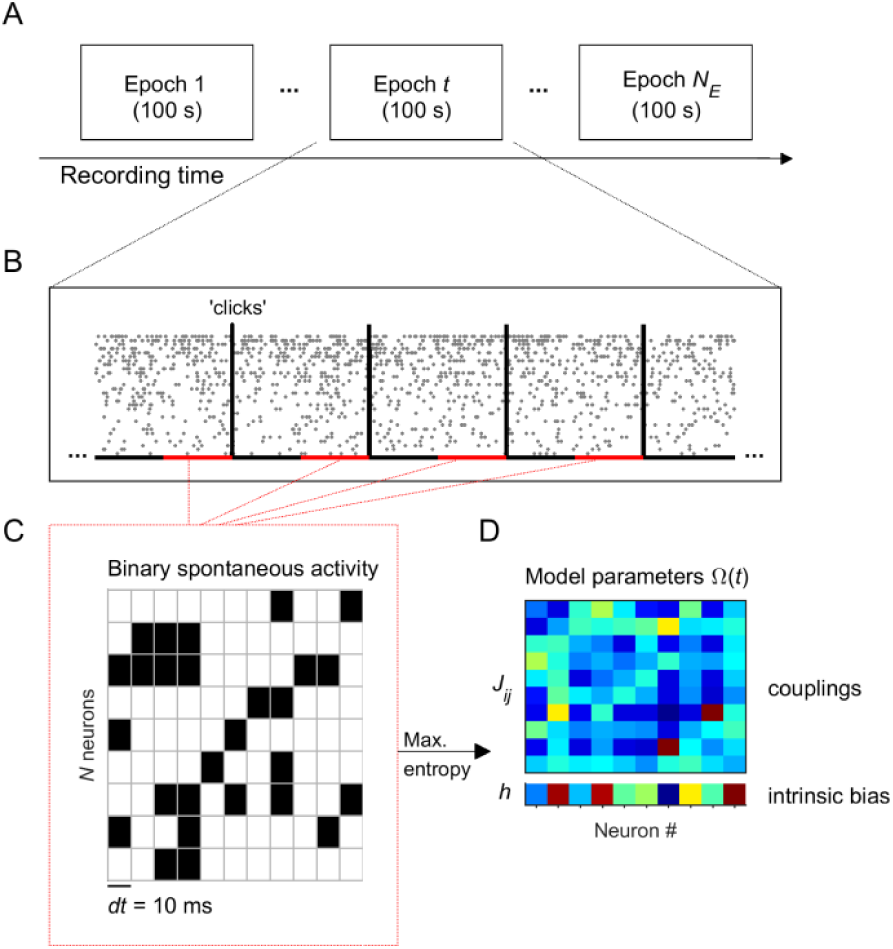
Experiment and analysis designs. **A:** Each recording session was divided into *N*_*E*_ adjacent epochs of 100 s. **B:** Each epoch contained a series of stimulus presentations. Stimuli consisted on acoustic clicks. For each 100-s epoch we collected the spontaneous activity, i.e., the activity during 1.5-s intervals preceding each stimulus (red intervals), to build concatenated binary data. **C:** Binary data was obtained by discretizing time in bins of *dt* = 10 ms. Within each time bin, the ensemble activity of *N* neurons was described by a binary vector, 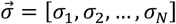, where *σ*_*i*_ = +1 if the *i*-th neuron generated a spike (black) and *σ*_*i*_ = −1 otherwise (white). **D:** Maximum entropy models were used to describe the binary patterns of subsets of 10 neurons, during each 100-s epoch. The model parameters Ω = {*h, J*} represents the intrinsic tendency of neuron *i* towards activation (*σ*_*i*_ = +1) or silence (*σ*_*i*_ = −1), noted *h*_*i*_, and the effective interaction between neurons *i* and *j*, noted *J*_*ij*_.

### Description of spontaneous activity patterns using maximum entropy models

We first examined the temporal evolution of the spontaneous activity across the *N*_*E*_ epochs. Because we were interested in the evolution of the statistics of ensemble activity, we described the collective activity of groups of *N* single-units using a maximum entropy model (MEM) (Schneidman et al., 2006; Shlens et al., 2009; Tkačik et al., 2015) in each epoch (see **Methods** and **Figure 1C–D**). These models allowed us to describe the patterned activity with a small number of parameters. To fit the model, time was discretized in bins of *dt* = 10 ms. Within each time bin, the ensemble activity of *N* neurons was described by a binary vector, 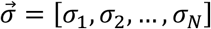, where *σ*_*i*_ = +1 if the *i*-th neuron fired a spike in that time bin and *σ*_*i*_ = −1 otherwise. The collective activity was determined by the probability distribution 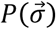 over all 2^*N*^ possible binary patterns. The MEM fits 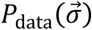 by finding a distribution 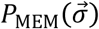 that maximizes its entropy under the constraint that the activation rates (< *σ*_*i*_ >) and the pairwise correlations (< *σ*_*i*_*σ*_*j*_ >) found in the data are preserved in the model. It is known that the maximum entropy distribution that is consistent with these constraints is the Boltzmann distribution, 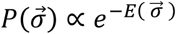, where 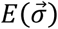 is the energy of the pattern 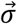, given by: 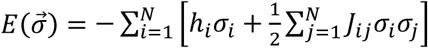 (Schneidman et al., 2006; Tkačik et al., 2015). The model parameter *h*_*i*_ represents the intrinsic tendency of neuron *i* towards activation (*σ*_*i*_ = +1) or silence (*σ*_*i*_ = −1) and the parameter *J*_*ij*_ represents the effective interaction between neurons *i* and *j*. Once we learned the parameters **Ω** using a gradient descent algorithm (see **Methods**), the expected probability of any pattern is known. For each recording session, we fitted the model using the spontaneous binarized activity from an ensemble of *N* = 10 randomly selected single neurons separately for each of the *N*_*E*_ epochs. We chose *N* = 10 because 100-s epochs provided around 5000 observed spontaneous patterns, which is a reasonable amount to get an estimate of the distribution of the 2^10^ = 1024 possible patterns. To accurately estimate models of larger *N*, the epochs ought to be much larger preventing possibility to investigate the temporal evolution of the model along the experiment. We finally repeated the process of randomly choosing *N* = 10 single units *Q* times for each experiment (for datasets 3 and 5: *Q* = 10 ensembles, otherwise: *Q* = 20). In summary, for each recording session, we built *Q* × *N*_*E*_ models, each composed of 10 units. Before studying the evolution of the model parameters **Ω**(*t*) across epochs (*t* = 0, 1, 2..), we first evaluated how well the MEM fitted the data.

For each epoch, we used the Jensen-Shannon divergence (*D*_*JS*_, see **Methods**) to measure the similarity between the probability distribution of the empirical and model binary patterns (**Figure 2A-C**). We compared this similarity to the distribution of binary patterns predicted from independent-MEMs, for which only the activation rates were preserved (i.e., only ***h*** was optimized). We found that the empirical distribution was well approximated by MEMs and that, for all recording sessions, the goodness-of-fit (i.e., 1/*D*_*JS*_) was orders of magnitude higher for MEMs than for independent-MEMs (**Figure 2C**), leading to excellent model performances (i.e., Kullback-Leibler ratio equal to 0.95 ± 0.03 on average, see **Methods**).

**Figure 2.**
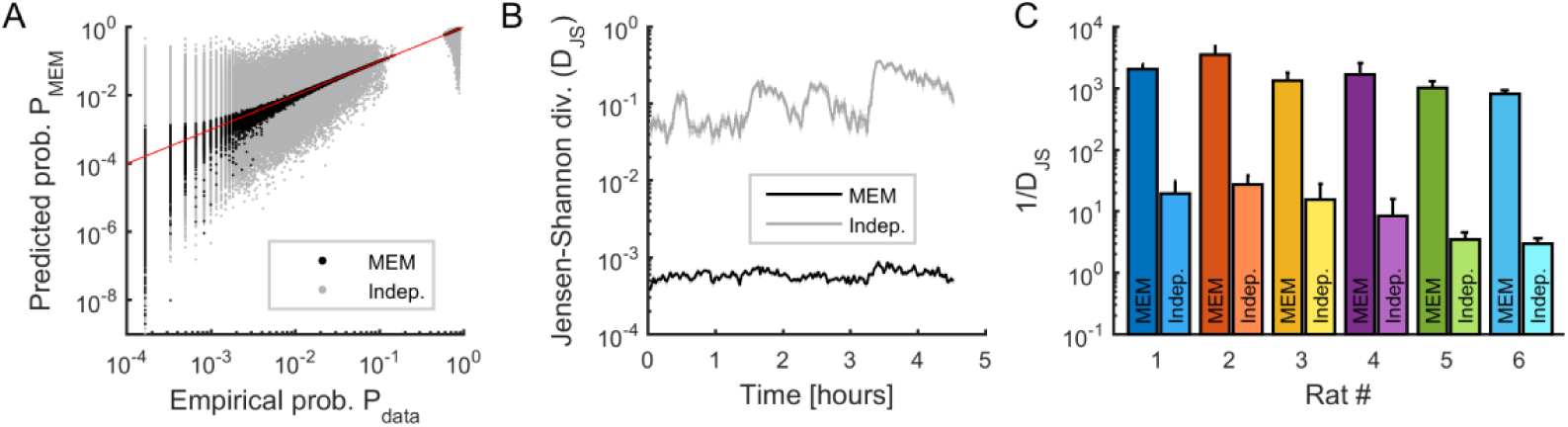
Fitting maximum entropy models (MEMs) to spontaneous activity patterns. **A:** Comparison between the probability distribution of empirical binary patterns and the probability distribution predicted by MEMs (black dots) and independent models (gray dots), for all epochs. Every point represents a binary pattern that has appeared in the data at least once. Red line represents the identity line. **B:** Jensen-Shannon divergence (D_JS_) between spiking data and MEMs, and between spiking data and independent models, across time, averaged across neuronal ensembles. Data in (A) and (B) correspond to one example rat (#1). **C:** Averaged goodness-of-fit (1/D_JS_) for MEMs and for independent models, for all rats. Error bars indicate SEM.

### Temporal evolution of activity observables, model parameters, and their sensitivity

We next analyzed the temporal evolution of the different spiking data statistics and the model parameters. We first measured the temporal variation of the activity observables (i.e., firing rates and pairwise correlations) by calculating the average Pearson correlation (or similarity *γ*; see **Methods**) between the values in epoch *t* and those in epoch *t* + Δ*t* (**Figure 3A**). This similarity rapidly decayed with Δ*t*, indicating that the observables substantially changed over time. We next examined how much these variations influenced the evolution of the collective activity characterized by the distribution of binary patterns. For this, we evaluated how well the data in a given epoch *t* could be explained by the MEM constructed using the data at time *t* + Δ*t.* Specifically, we calculated ⟨*D*_*JS*_⟩(Δ*t*), given by the average Jensen-Shannon divergence between the distribution of data binary patterns in epoch *t*, i.e. *P*_data, *t*_, and the distribution of binary patterns predicted by the MEM constructed using the data in epoch *t* + Δ*t*, i.e.*P*_*MEM, t* + Δ*t*_ (see **Methods**). We found that ⟨*D*_*JS*_⟩(Δ*t*) increased as a function of Δ*t*, indicating that the model parameters changed during the recording session, leading to changes in collective activity (**Figure 3B**). Indeed, the similarity of the model parameters also decayed rapidly over time (**Figure 3C**, *orange* and *purple* traces).

**Figure 3.**
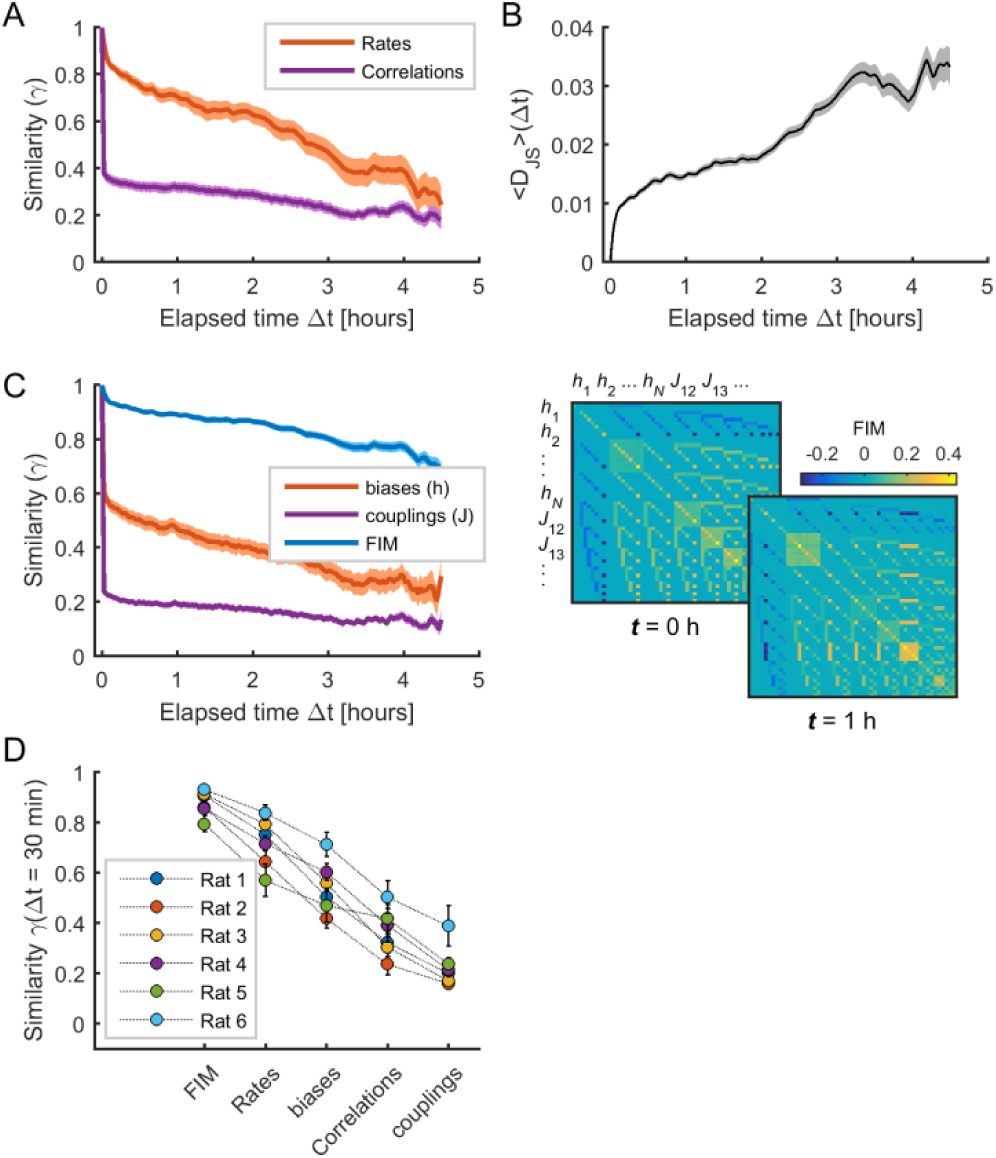
The sensitivity of model parameters was more stable than activity observables. **A:** Similarity (i.e., Pearson correlation coefficient) of mean firing rates (red) and pairwise correlations (purple) as a function of elapsed time Δ*t.* **B:** Jensen-Shannon divergence (D_JS_) between the distribution of empirical spiking patterns in epoch *t* and the distribution of binary patterns of the pairwise MEM in epoch *t* + Δ*t*, averaged over all *t*. **C:** *Left*: Similarity of Fisher information matrix (FIM) elements, biases (*h*_*i*_), and couplings (*J*_*ij*_) as a function of elapsed time Δ*t. Right:* FIMs at time *t* = 0h and *t* = 1h. Data in (A), (B), and (C) correspond to one example rat (#1); shaded areas correspond to SEM. **D:** Similarity of FIM elements, rates, biases, correlations, and couplings after 1/2 hour (i.e., Δ*t* = 30 min), for all rats. Error bars indicate SEM.

As shown in Panas et al. (2015), changes in model parameters can differently contribute to collective activity, since the model can be sensitive to changes in some few combinations of parameters. Following this, we next evaluated the sensitivity of model parameters by calculating the Fisher information matrix (FIM, see **Methods**) for each neuronal ensemble and each epoch. The FIM measures how much the model log-likelihood 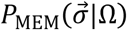 changes with respect to changes in the parameters **Ω**. We first notice that the FIM had the highest stability across time, compared to the data firing rates and correlations and the model parameters (**Figure 3C**, *blue* trace). Indeed, the similarity after 1/2 hour was *γ* = 0.882 ± 0.002 for the FIM, *γ* = 0.732 ± 0.003 for the firing rates, *γ* = 0.551 ± 0.004 for the biases, *γ* = 0.364 ± 0.004 for the correlations, *γ* = 0.234 ± 0.003 for the couplings (*F*_4,495_ = 305.73, p<0.001, one-way ANOVA followed by Tukey’s post hoc analysis) (**Figure 3D**). Altogether these results show that the sensitivity of the model parameters remained relatively stable despite substantial changes in firing rates, correlations, collective activity and the model parameters themselves.

### Spontaneous neuronal activity presents sloppiness

Having shown that the sensitivity of model parameters was relatively stable during the recording sessions, we next studied the structure of the FIMs. First, we noted that most elements of the FIM had near-zero values (**Figure 4A**) indicating that most of the parameters had a small effect on the model log-likelihood. In contrast, a small fraction of elements had values strongly different from zero as revealed by the heavy tail of the distribution of FIM values (**Figure 4A**). To identify the parameter combinations that had the strongest effect on model behavior, we decomposed the FIM into eigenvectors and classified them according to their eigenvalue (**Figure 4B**). We observed that, except for some few eigenvalues, most of the FIM eigenvalues were small, corresponding to combinations of parameters that had little effect on model behavior. These unimportant parameter combinations defined the sloppy dimensions of the model. The few eigenvectors with large eigenvalues defined the stiff parameter dimensions along which the model behavior was strongly affected.

**Figure 4.**
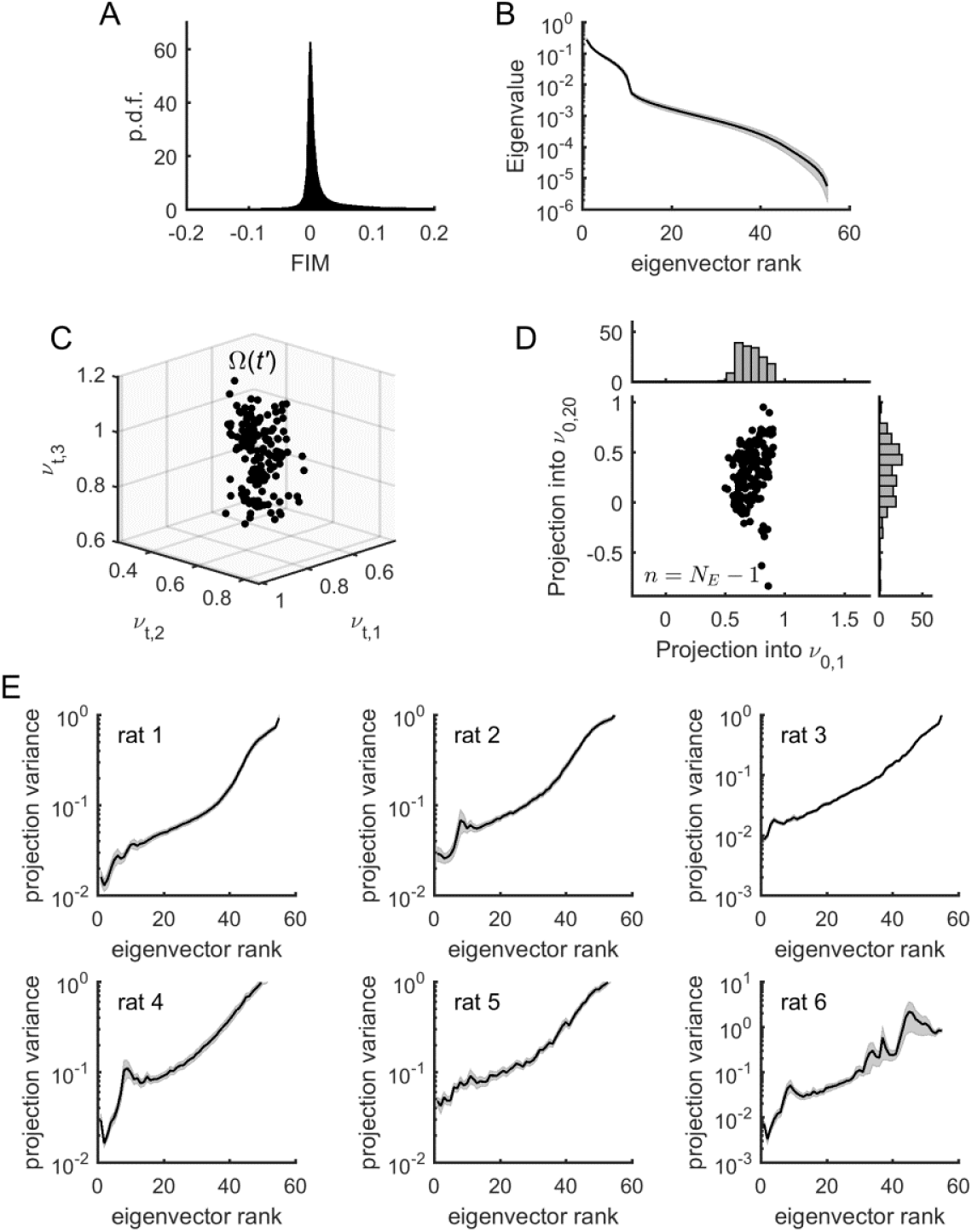
Model parameters presented sloppiness and predominantly evolved along sloppy dimensions. **A:** Distribution of FIM elements, for all epochs and all rats. **B:** Eigenvalues of the FIM, average across all epochs, for an example rat (# 1). Shaded areas represent SEM. **C:** Projection of Ω(*t*′) into the first three eigenvectors of the FIM from a given epoch *t*, for all *t*′ ≠ *t*. Data from rat 1. **D:** Projection of Ω(*t*′) into the first and the 20th eigenvectors of the FIM from epoch *t*, noted *v*_*t*,1_ and *v*_*t*,20,_ respectively, for all *t*′ ≠ *t. Top inset*: distribution of projections into *v*_*t*,1_. *Right inset*: distribution of projections into *v*_*t*,20_. Note higher variance of projections into *v*_*t*,20_ than into *v*_*t*,1_. Data from rat 1. **E:** Average variance of projections of Ω(*t*′)into the different eigenvectors of the FIM at epoch *t* (for all *t*′ ≠ *t*), for the different rats. Shaded areas represent SEM.

In the following we showed that the temporal evolution of the model parameters occurred predominantly along the sloppy dimensions. For this, we projected the parameters **Ω**(*t*′), calculated at time *t*′, into the eigenvectors of the FIM at time *t*, denoted ***v***_*t*,1_, ***v***_*t*,2_, …, ***v***_*t,k*_, …, where *k* is the rank of the eigenvector (**Figure 4C**). For each dimension, or eigenvector, we obtained a distribution of projections of parameters **Ω**(*t*′) (**Figure 4D**). To quantify how much the parameters varied along each eigenvector, we calculated the average variance of each projection as a function of the rank of the eigenvector. We found that the projection variance increased as a function of the eigenvector’s rank for all datasets (**Figure 4E**). This indicates that the model parameters predominantly evolved along sloppy dimensions (i.e., FIM eigenvectors of highest rank *k*), while they remained relatively stable along stiff dimensions (i.e., FIM eigenvectors of lowest rank *k*). Using stationary surrogate data, we controlled that these parameter fluctuations were not fully explained by estimation errors and, furthermore, that parameter fluctuations along sloppy dimensions were those that deviated the most from the stationary case (**Figure S1**). Nevertheless, we noted that the projection variance into the stiff dimensions, albeit small, was not zero. This means that the model also evolved along parameter dimensions that had a strong impact on the collective activity. We hypothesized that changes in collective behavior, associated to changes in stiff parameters, were related to changes in cortical state.

### Cortical state transitions evolve along stiff dimensions

To test this hypothesis, we first measured the cortical state in each epoch *t* using silence density, *CS*(*t*), defined as the fraction of 20-ms time bins with zero population activity, i.e. no spikes from any neuron (see **Methods**) (Luczak et al., 2013; Pachitariu et al., 2015; Mochol et al., 2015). To obtain the most accurate estimate of silence density, we used all the spikes from the merge of all the single-units and multi-units in the calculation of *CS*(*t*). During the course of the experiment, we observed large fluctuations in silence density, with low and high values associated to de-synchronized and synchronized cortical states, respectively (**Figure 5A**). We found that differences in collective dynamics in different epochs, quantified by *D*_*JS*_ (*t,t′*) = *D_JS_*(*P*_data, *t*_;*P*_MPM, t′_), significantly co-varied with the changes in cortical state, given by *d* = |*CS*(*t*′) – *CS*(*t*)| (averaged correlation coefficient 0.60 ± 0.08, p < 0.001) (**Figure 5B-C**). Thus, changes in collective behavior correlated with changes in cortical state.

**Figure 5.**
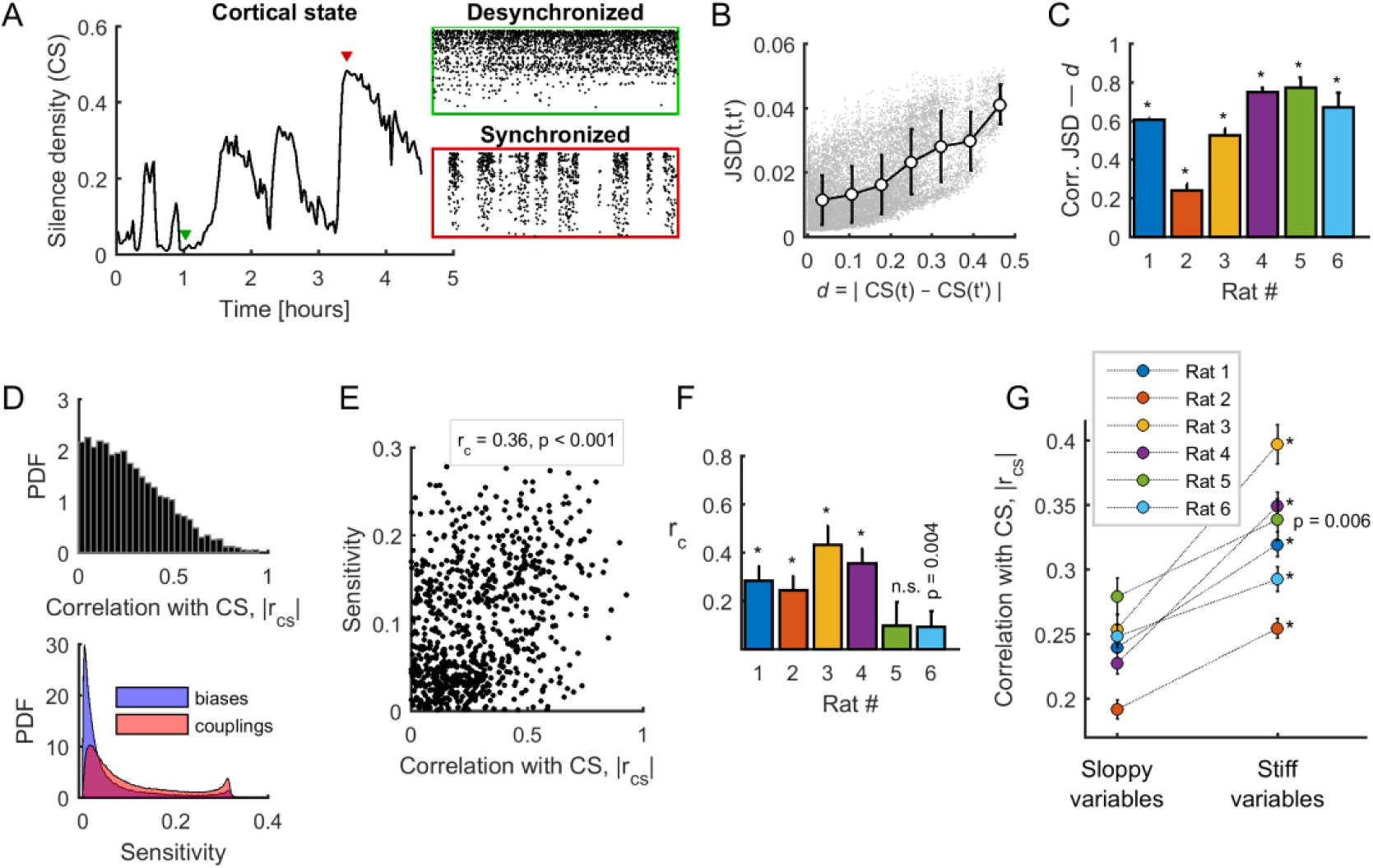
Spontaneous neuronal activity evolved along stiff dimensions during cortical state transitions. **A:** Silence density was used to characterize the cortical state. *Green inset*: high values of the silence density indicate desynchronized cortical states (each row represents the spike train of a single-unit). *Red inset*: low values of the silence density indicate synchronized cortical states. Data from rat 1. **B:** Difference in collective pattern statistics, i.e. *D*_*JS*_, between different epochs, *t* and *t′*, as a function of the corresponding difference in silence density, noted *d*. Each gray dot corresponds to a pair of epochs (*t, t′*). The solid line indicates the average relation between *D*_*JS*_ and *d*; error bars indicate SD. Data from rat 1. **C:** Correlation coefficient between *D*_*JS*_ and *d*, for all rats. *: p < 0.001. **D:** *Top*: Distribution of the absolute value of the correlations between the cortical state and the activity observables, noted |*r*_*cs*_|. *Bottom*: Distribution of parameter sensitivity values for biases (*h* parameters) and couplings (*J* parameters), for all models. **E:** |*r*_*cs*_| vs. sensitivity. Correlation: *r*_*c*_ = 0.36; p <0.001. Data from rat 4. **F:** Correlation between |*r*_*cs*_| and the sensitivity of all activity variables (i.e., firing rates and pairwise correlations), for each dataset. *: p < 0.001. Error bars indicate correlation 95% confidence interval. **G:** |*r*_*cs*_| for sloppy and stiff variables. *: p < 0.001, paired *t*-test. Error bars indicate SEM.

We next asked which activity observables, i.e., the firing rate of each neuron and all pairwise correlations, related more to cortical state transitions. For this, we calculated the absolute correlation, |*r*_*cs*_|, between the cortical state *CS*(*t*) and the activity observables. We found that |*r*_*cs*_| was broadly distributed between 0 and 0.94, thus some observables correlated more with the cortical state (**Figure 5D**, top panel). Next, to relate the sensitivity of model parameters (their stiffness) to the activity observables, we measured the sensitivity of a given parameter by its average contribution to the first eigenvector of the FIM, i.e., the sensitivity of *i*-th parameter is given by 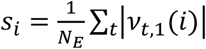, and we associated it to the corresponding observable. Note that the ranges of the sensitivity of biases (*h*) and couplings (*J*) were similar (**Figure 5D**, bottom panel), and that sensitivities calculated in the first and the second halves of the recording session were highly correlated (correlation coefficient > 0.82, for all rats; average: 0.89 ± 0.03). We found a significant positive correlation between the associated sensitivity (*s*) and the correlation with the cortical state (|*r*_*cs*_|) in 5/6 datasets (**Figure 5E-F**). Thus, the observables that correlated more with the cortical state were those with the highest associated sensitivity. This result led us to separate the activity observables into two classes, called “sloppy” and “stiff”, based on whether the associated sensitivity (*s*) was lower or higher than the median *s*. We found that stiff variables were significantly more correlated with the cortical state than the sloppy variables (p<0.01 for all datasets, paired *t*-test; **Figure 5G**). Altogether, these results indicate that neuronal activity and co-activity preferentially evolved along sensitive (stiff) parameter dimensions during cortical state transitions.

### Sensory-evoked activity evolves along sloppy dimensions

The above results indicate that, although intrinsic spontaneous dynamics predominantly evolved along sloppy dimensions (**Figure 4F**), cortical state transitions were governed by changes in stiff parameters (**Figure 5G**). We next investigated which parameter dimensions were explored when the neural network was driven by external sensory inputs, i.e. during stimulus-evoked activity (**Figure 6A**). We observed that evoked responses (which could be increased or decreased with respect to pre-stimulus baseline firing rate) were larger for sloppy neurons than for stiff neurons (**Figure 6B-C**). To quantify the responsiveness of each neuron, we calculated the modulation index (MI, see **Methods**) of each neuron in response to acoustic stimuli. We next calculated the relation between MI, calculated during evoked activity, and the sensitivity *s* associated to firing rates, calculated during the spontaneous activity as above. We found that the more responsive neurons were those with the lowest associated sensitivity (**Figure 6D-E**). This indicates that stimulus-evoked neuronal activity evolved mostly along sloppy dimensions. Finally, we evaluated the difference, noted ΔMI, between the MI of sloppy and stiff neurons as a function of cortical state *CS*(*t*). Specifically, first, the MI values in each epoch were averaged according to different ranges of the silence density. Second the MI values of sloppy and stiff neurons were compared within each range. We found that ΔMI was maximal during desynchronized activity, and minimal during synchronized activity (**Figure 6F**). Thus, the cortical activity during stimulus response evolved predominantly along sloppy dimensions for the desynchronized cortical state, while, in the synchronized state, the dominance of sloppy fluctuations was reduced, and stiff fluctuations became comparable.

**Figure 6.**
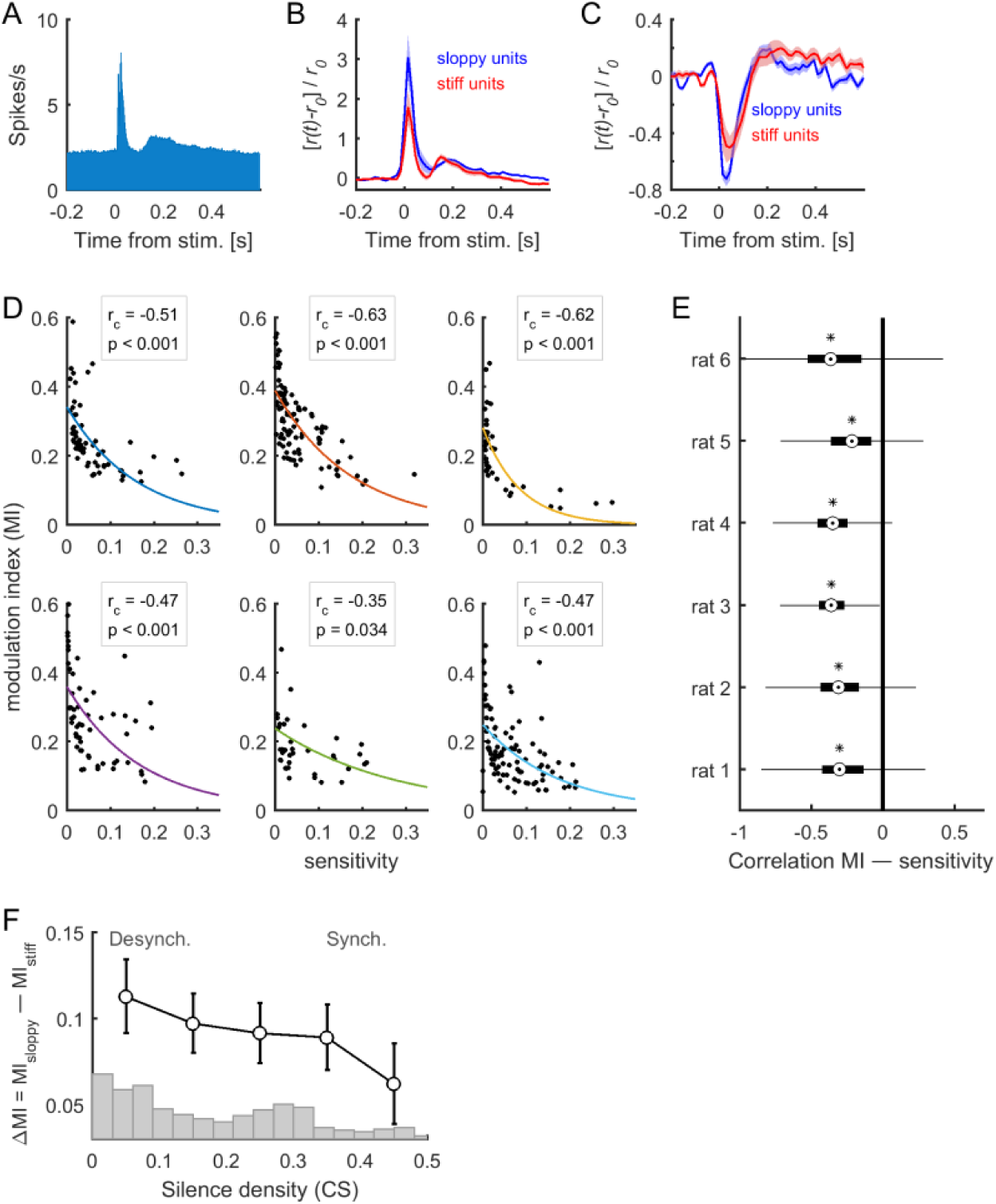
Stimulus-evoked neuronal activity evolved along sloppy dimensions. **A:** Population responses to acoustic clicks. **B-C:** Median-split of sensitivity was used to separate stiff neurons and sloppy neurons. The mean responses for stiff and sloppy neurons are shown in the case of excited responses (B) and suppressed responses (C). The responses were normalized by the average pre-stimulus activity *r*_0_. Shaded areas correspond to SEM. Data in (A), (B), and (C) correspond to one example rat (#1). **D:** Modulation index (MI) as a function of associated sensitivity of firing rates (*s*), for each dataset. The correlation between MI and *s* was negative for all datasets (*r*_*c*_: correlation coefficient; p: p-value). Solid lines indicate exponential fits. **E:** Correlation between MI, calculated in epoch *t*, and ***v***_*t*,1_, for all rats. On each box, the central mark indicates the median, and the bottom and top edges of the box indicate the 25th and 75th percentiles, respectively. Asterisks indicate significantly negative medians (p < 0.001, two-sided signed rank test). **F:** Difference of the MI of sloppy neurons minus the MI of stiff neurons as a function of cortical state (i.e., silence density), averaged for all rats (black trace; error bars indicate SEM). The gray bars indicate the distribution of silence density values.

### Stiff parameters were associated to central neurons within the neuronal network

In this section, we further investigate the properties of neurons and pairs of neurons with respect to their associated parameter sensitivity. As above, we separated the neurons and pairs of neurons into two classes, called “sloppy units/pairs” and “stiff units/pairs”, based on whether the associated sensitivity (*s*) was lower or higher than the median *s* (units were associated to parameters *h*_*i*_, and pairs or links were associated to parameters *J*_*ij*_). We first found that stiff units were significantly more active than sloppy units (**Figure 7A**). We quantified this by performing receiver operating characteristic (ROC) analysis and used the area under the ROC curve (AUC) as a measure of how well the firing rates distributions of the two classes were separated (AUC = 0.961–0.998, p < 0.001, for all rats; **Figure 7F**). Stiff neurons were also significantly more correlated among them than sloppy neurons (AUC = 0.615–0.932, p < 0.001, for all rats; **Figure 7B**,**F**). The distributions of correlations remained well separated when calculated for the links, i.e, pairs of neurons with associated parameters *J*_*ij*_ (AUC = 0.541–0.766, p < 0.001, for all rats; **Figure 7C,F**).

**Figure 7.**
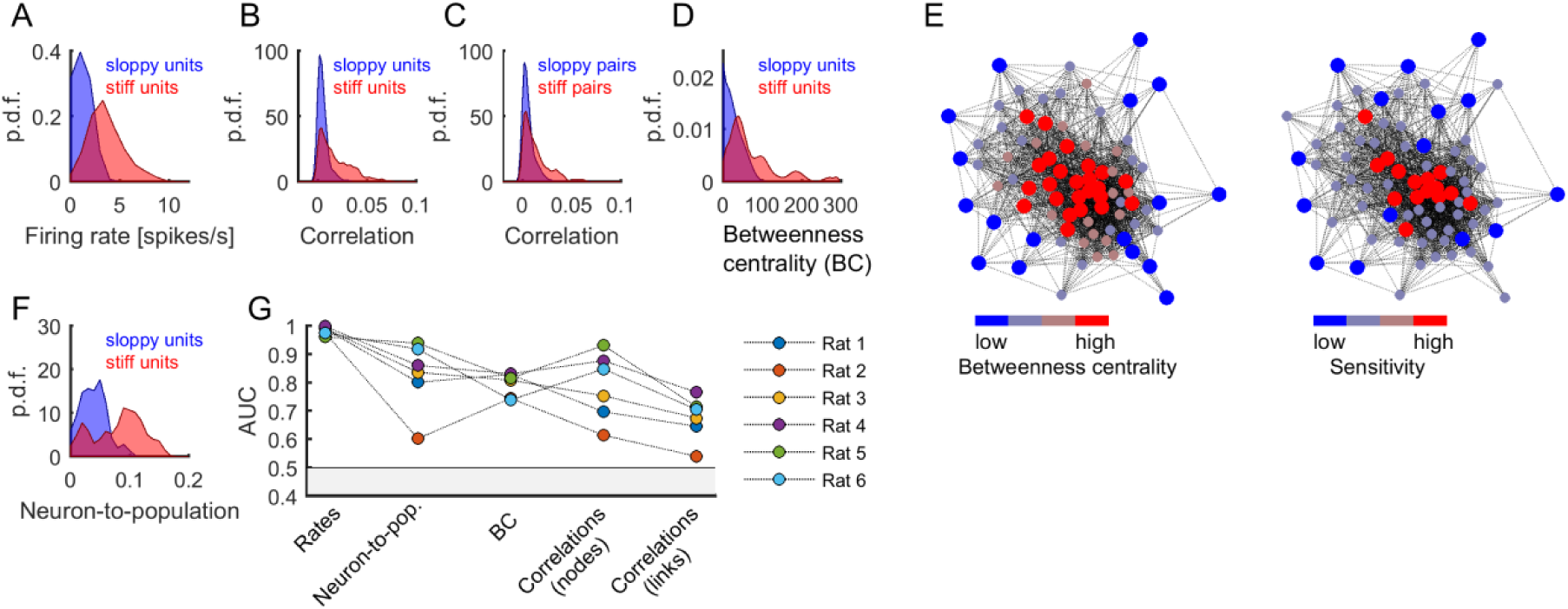
Central and highly active neurons were associated to stiff parameters. **A:** Distribution of firing rates of sloppy and stiff neurons. **B:** Distribution of correlations among sloppy neurons and among stiff neurons. **C:** The distribution of correlations was also calculated for the links (i.e, pairs of neurons with associated parameters *J*_*ij*_). Note that, in principle, links can be related to pairs composed of one sloppy and one stiff neuron. **D:** Distribution of betweenness centrality of sloppy and stiff neurons. **E:** Connectivity graph: each node represents a neuron and links represent significant correlations between pairs of neurons. The graph was plotted using force-directed layout, i.e. using attractive forces between strongly connected nodes and repulsive forces between weakly connected nodes. *Left*: the nodes were colored as a function of betweenness centrality. *Right*: the nodes were colored as a function of associated sensitivity. Note the high overlap between both color labeling methods, indicating that sensitivity was highly predictive of the centrality of the nodes. **F:** Distribution of neuron-to-population couplings of sloppy and stiff units. Panels A–F show data from rat 1. **G:** Area under the receiver operating curve (AUC) quantifying the separation of distributions of sloppy and stiff classes. All AUC values were significantly higher than 0.5 (p < 0.001).

To further investigate the structure of correlations, we evaluated the centrality of stiff and sloppy neurons within the observed network of neurons. For this we used the betweenness centrality (BC), a measure of node centrality in a graph or network, which in our case was given by the functional connectivity matrix among the recorded neurons (see **Methods**). The BC measures the extent to which a node in the graph tends to lay on the shortest path between other nodes. Thus, a node with higher BC has more influence over the network, because more information passes through that node. We found that stiff neurons had significantly more centrality in the functional connectivity graph than sloppy neurons (AUC = 0.740–0.831, p < 0.001, for all rats; **Figure 7D,F**). This indicates that stiff neurons were part of the core of the graph, while sloppy neurons were part of the graph periphery, as clearly shown using graph visualization (**Figure 7E**) (Fruchterman and Reingold, 1991). BC values were correlated with firing rates (correlation coefficient: 0.59 ± 0.11), which could suggest that differences in BC between stiff and sloppy neurons were simply a consequence of differences in firing rates. However, using surrogate data that preserved the observed firing rates and produced correlations through global modulations, we found that neither the structure of correlations nor the BC values could be trivially predicted by globally modulated firing rates but they were rather suggestive of functional interactions (**Figure S2**). Thus, in addition to different firing rates, different correlations and BC values were supplementary features of stiff and sloppy neurons.

Moreover, previous work has shown that cortical neurons differ in their coupling to the population activity, with neurons that activate most often when many others are active and neurons that tend to activate more frequently when others are silent (Okun et al., 2015). Thus, along with centrality, we calculated the neuron-to-population coupling, given by the Pearson correlation between the activity of each neuron *i* and the number of coactive neurons (excluding neuron *i*; see **Methods**). We found that stiff neurons were significantly more coupled to the population activity than sloppy neurons (AUC = 0.603–0.939, p < 0.001, for all rats; **Figure 7D,F**). In summary, stiff units were more active, more central, more coupled among them, and more coupled to the population activity than sloppy units.

## Discussion

We here studied the changes in activity caused by intrinsic (i.e. cortical state) and extrinsic (i.e., stimulus-evoked) sources in A1 neuronal ensembles in an estimated parameter space. The parameter space was obtained using the maximum entropy principle, providing a handful number of parameters describing the probability of all possible binary activity patterns. These parameters differed in their impact on collective activity that was sensitive to a few combinations of parameters, called stiff dimensions, but insensitive to many others called sloppy dimensions. Our results suggest that spontaneous cortical state transitions and stimulus-driven activity evolved along different parameter dimensions. Indeed, in one hand, while most of the fluctuations during spontaneous activity evolved along sloppy dimensions, some residual ongoing fluctuations evolved along stiff dimensions, and these fluctuations were correlated with synchronized/desynchronized cortical state transitions. On the other hand, stimulus-induced activity was larger in sloppy dimensions than in stiff dimensions, an effect that was most prominent during the desynchronized cortical state. Note that the observation that both spontaneous and stimulus-driven activities predominantly evolve along sloppy dimensions results from the strong similarity of spontaneous and evoked activity, reported in several previous studies (Arieli et al., 1996; Kenet et al., 2003; MacLean et al., 2005; Luczak et al., 2009). Finally, by classifying the neurons as stiff versus sloppy neurons (i.e., those contributing more or less to the principal stiff dimension) we found that the firing rates and the functional connectivity topology significantly differed between the two classes of neurons.

The observation that stimulus-induced activity evolved along sloppy dimensions can have important functional implications. It suggests that a stimulus can modulate the activity of a subset of sloppy neurons without entirely affecting the collective activity. This could be an efficient functional architecture to encode sensory information without perturbing other ongoing or memory-stored processes. Consistent with this view and with previous studies (Margolis et al., 2012; Panas et al., 2015), our results suggest that the integrity of the network is ensured by a core of highly active stiff neurons, which have strong functional connections among them (either through anatomical connections or common inputs), topologically peripheral sloppy neurons (within the functional connectivity graph) can be largely modulated by external inputs. A similar sub-network of highly active, interconnected neurons has been recently identified in the mice neocortex (Yassin et al. 2010). Importantly, sensory input was not required to drive these cells. Previous studies of complex systems have derived general principles of core/periphery network structures: the network periphery is more variable, evolvable, and plastic than the network core, while the network core facilitates system robustness (Kitano, 2004; Csermely et al., 2013). Thus, we hypothesize that sloppy neurons could also be more affected by synaptic plasticity, allowing for network reconfiguration without loss of stability. Consistent with this, previous work on whole-brain fMRI has observed core stability and peripheral flexibility over the course of learning (Bassett et al., 2013). Furthermore, we observed that stimulus responses evolved more pronouncedly along sloppy dimensions in the desynchronized state, while in the synchronized state fluctuations along sloppy and stiff dimensions were comparable (**Figure 6F**). This supports the view that responses along sloppy dimensions provide information processing benefits, since previous studies have shown that auditory stimuli in rodents (Marguet and Harris, 2011; Pachitariu et al., 2015) and visual stimuli in both rats (Goard and Dan, 2009) and monkeys (Beaman et al., 2017) are better represented in the desynchronized state as compared to the synchronized state. Finally, the properties of spontaneous and induced cortical dynamics observed in the present anesthetized condition are likely to be relevant also during wakefulness. Indeed, several studies reported the existence of synchronized cortical states during wakefulness (for review see Zagha and McCormick, 2014), and global fluctuation resembling transitions between up and down periods during alert or quiescent wakefulness (Petersen et al., 2003; Luczak et al., 2007; Poulet and Petersen, 2008; Zagha et al., 2013; Tan et al., 2014; Engel et al. 2016) or even during task engagement (Sachidhanandam et al. 2013).

We found that stiff neurons were more linked to the observed neuronal population activity than sloppy neurons. Stiff neurons had higher centrality in the functional connectivity graph and higher coupling to the population activity than sloppy neurons. Stiff neurons could thus sense and influence larger parts of the network. Previous research showed that neurons differ in their coupling to the population activity, with neurons that activate most often when many others are active, called “choristers”, and neurons that tend to activate more frequently when others are silent, called “soloists” (Okun et al., 2015). Our results suggest that stiff and sloppy neurons are chorister and soloist neurons, respectively. In other words, changes in the activity of stiff/chorister neurons lead to changes in collective behavior (i.e., cortical states), while the activity of sloppy/soloist neurons can spontaneously fluctuate or respond to stimuli without strongly affecting the collective behavior. Thus, we believe that the roles of stiff/chorister neurons and sloppy/soloist neurons are important to understand tradeoffs between responsiveness and stability of the network. Furthermore, we here studied the evolution of neuronal activity on the time scale of hours and found that fluctuations on stiff parameter dimensions were the weakest and were related to cortical state transitions, which time scale is in the order of tens of minutes (Hahn et al., 2017; Mochol et al., 2015). Previous studies have reported prominent changes on neuronal activity and tuning properties over days, but with stable decoding performances of population activity (Chestek et al., 2007; Ziv et al., 2013; Panas et al., 2015). However, we hypothesize that learning or adaptation to changing environments could lead to large changes in collective activity. In that case, particular attention could be paid to the influence of high-order areas on the activity of subsets of stiff and sloppy neurons from sensory areas, as top-down regulation might be a mechanism to control the stabilizing network core.

The coexistence of two subgroups of cortical neurons (stiff and sloppy) provides new valuable architectural constrains for models of the brain state and its transitions. Several past studies have modelled the synchronized brain dynamics as transitions between two attractors. Depending on the model specificity those transitions could be noise driven (Mejias et al., 2010; Mochol et al., 2015, Jercog et al.,2017) or caused by some fatigue mechanism (Compte et al., 2003; Hill and Tononi, 2005; Mattia and Sanchez-Vives, 2012). To make the system works in a desynchronized regime it was enough to increase the background input to the network (Bazhenov et al., 2002; Hill and Tononi, 2005; Curto et al., 2009; Destexhe, 2009; Mochol et al., 2015). Given our present results, the models could be extended to include a network core/periphery architecture, a non-homogeneous background input preferentially targeting the network core, and different stimulus spatial distributions. Such a model would provide insights on the interplay between cortical state transitions and sensory representation. Moreover, our findings question the view that the mechanisms by which background and stimulus inputs impact the dynamics are similar, as assumed in the simple bi-stable rate model (Mochol et a. 2015).

Finally, we here described the patterned activity of small (*N* = 10) neuronal ensembles using MEMs. It is known that MEMs of small sizes can present departures from the observed distribution of summed activities and higher-order correlations (Tkačik et al., 2014). Recent advancements on learning algorithms allow to construct MEMs of ∼100 neurons. However, these models cannot be used in a time-resolved manner, as we did here, due to limited data in each epoch. Small model sizes are thus the cost to pay to study the evolution of collective activity over time in a meaningful time scale (i.e., the one of cortical state transitions).

## Methods

### Experimental Techniques

We analyzed the neuronal activity recorded in the primary auditory cortex (A1) of 6 anesthetized rats (Sprague–Dawley; 250–400 g). The experimental procedures and spikes sorting procedures have been previously described in Mochol et al. (2015). Briefly, rats were anesthetized with urethane (1.5 g/kg body weight) and silicon microelectrodes (Neuronexus) with 32 or 64 channels were inserted in deep layers (depth, 600–1,200 μm) of the primary auditory cortex. The spiking activity from single units and multi-units (i.e., neurons that were not well isolated) was simultaneously recorded during spontaneous activity and in response to acoustic “clicks” (5-ms square pulses; interstimulus interval, 2.5 or 3.5 s; see **Table 1**). In some datasets, double clicks (5-ms square pulses; 50- or 100-ms inter-click interval) were also presented, but, in the present study, we analyzed only the responses to single click.

**Table 1.** Number of neurons (SU: single-units, MU: multi-unit), number of 100-s epochs, number of stimulus presentations in 100-s epochs, and number of neuronal ensembles, for each dataset.

|  | No. of neurons | No. of 100-s epochs | Stimulus presentations in 100-s epochs | No. of neuronal ensembles (*Q*) |
| --- | --- | --- | --- | --- |
| **Rat 1** | SU: 81; MU: 3 | 163 | 12–14 | 20 |
| **Rat 2** | SU: 147; MU: 13 | 74 | 12–14 | 20 |
| **Rat 3** | SU: 44; MU: 30 | 70 | 12–20 | 10 |
| **Rat 4** | SU: 72; MU: 103 | 59 | 10–20 | 20 |
| **Rat 5** | SU: 58; MU: 39 | 29 | 28–29 | 10 |
| **Rat 6** | SU: 112; MU: 83 | 28 | 17–29 | 20 |

### Cortical state

Long continuous recordings (mean, ∼2 h) were divided into *N*_*E*_ 100-s epochs, and cortical state was estimated in each epoch based on spontaneous pooled population activity, i.e., the merge of single and multiunit spike trains during the 1.5-s intervals preceding each stimulus presentation. Cortical state was quantified using silence density defined as the fraction of 20-ms time bins with no population activity. Silent and active periods were obtained from the merge of consecutive empty and nonempty bins, respectively.

### Maximum entropy models

The spontaneous spiking activity of ensembles of *N* single neurons was studied using statistical modeling based on maximum entropy principle. The ensemble activity was binarized in non-overlapping time bins of *dt* = 10 ms, during which neuron *i* either did (*σ*_*i*_ = +1) or did not (*σ*_*i*_ = −1) generate one or more spikes. The state of the neural ensemble is described by a binary pattern 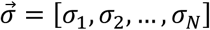, and thus the collective activity is described by the probability distribution 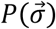 over all 2^*N*^ possible binary patterns. We estimated 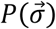 using a Maximum entropy model (MEM). The MEM finds 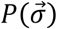 by maximizing its entropy under the constraint that some empirical statistics are preserved. A pairwise-MEM provides a solution under the constraint that the activation rates (< *σ*_*i*_ >) and the pairwise correlations (< *σ*_*i*_*σ*_*j*_ >) are preserved. The maximum entropy distribution 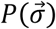 that is consistent with these expectation values is given by the Boltzmann distribution (Schneidman et al., 2006; Tkačik et al., 2015):

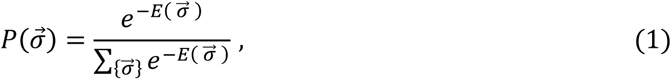

where 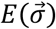 is the energy of the pattern 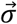, given by:

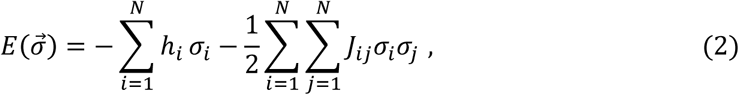

and 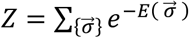 is the partition function.

The model parameter *h*_*i*_, called intrinsic bias, represents the intrinsic tendency of neuron *i* towards activation (*σ*_*i*_ = +1) or silence (*σ*_*i*_ = −1) and the parameter *J*_*ij*_ represents the effective interaction between neurons *i* and *j*. The estimation of the model parameters **Ω = {*h, J*}** was achieved through a gradient descent algorithm (see below). For each recording session, we constructed models for *Q* ensembles of *N* = 10 randomly selected single neurons and learned the model parameters using the spontaneous binarized activity within each 100-s epoch. Thus, for each recording session, we built *Q* × *N*_*E*_ models of 10 units. We were interested on the evolution of the model parameters over time, i.e., **Ω**(*t*) Note that number of free parameters is the sum of intrinsic biases and effective couplings, *N* + *N*(*N* − 1)/2 = 55, i.e., **Ω** = [*h*_1_, *h*_2_, …, *h*_*N*_, *J*_12_, *J*_13_, …].

### Estimation of MEM parameters

The MEM parameters **Ω = {*h, J*}**were iteratively adjusted to minimize the absolute difference between the empirical activation rates (⟨*σ*_*i*_⟩) and correlations (⟨*σ*_*i*_*σ*_*j*_⟩) and those (⟨*σ*_*i*_⟩_model_, ⟨*σ*_*i*_*σ*_*j*_⟩_model_) predicted by the model through Monte Carlo simulations. Specifically, each iteration is given by: 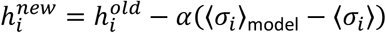, and 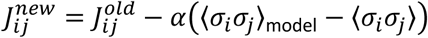, where *α* is the learning rate (*α* = 0.1). In our study we stopped the re-estimations once the differences between the empirical and model values are less than a tolerance threshold (0.005) or if this tolerance was not reached within a maximum number of iterations (100).

### MEM goodness-of-fit

The goodness-of-fit of the MEMs was evaluated using the Jensen– Shannon divergence (*D*_*JS*_) between the probability distribution of the empirical and model binary patterns (Marre et al., 2009). *D*_*JS*_ is a symmetric version of the Kullback-Leibler divergence (*D*_*KL*_) and is given as:

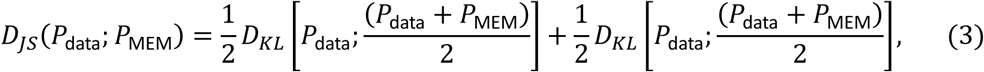

where:

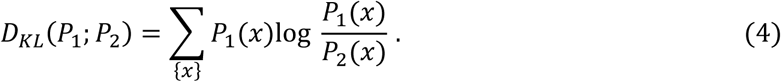

The fitting of MEM (second-order model) was compared to the fit obtained using independent-MEM, i.e., in which only for which only the activation rates (< *σ*_*i*_ >) are preserved (i.e., only ***h*** is optimized; first-order model). In this case, the pattern energy is given by: 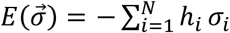.

Furthermore, the performance of the model can be evaluated using the Kullback-Leibler ratio, *R* (Shlens et al., 2009). This ratio is given by comparing the Kullback-Leibler divergence between the distribution *P*_1_ of the first-order model (i.e., independent-MEM) and the distribution of the actual data, *D*_1_ = *D*_*KL*_(*P*_1_;*P* _data_), with the Kullback-Leibler divergence between the distribution *P*_2_ of the second-order model and the distribution of the actual data, *D*_2_ = *D*_*KL*_(*P*_2_; *P*_data_). Specifically, the Kullback-Leibler ratio is defined as:

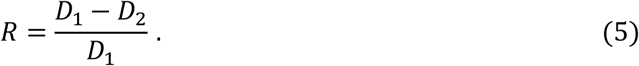

This ratio can range between 0 and 1, with 1 giving the highest performance.

### Fisher information matrix

Because in the MEM, all the information about the collective activity is contained in the probability distribution of the binary patterns, 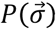, one can define the model parameter space as 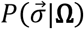 We were interested in knowing which parameters, or combination of parameters, have a strong effect on the collective activity. To measure how distinguishable two models, with parameters **Ω** and **Ω +** *δ***Ω**, are based on their predictions, we used the Fisher information matrix (FIM). Indeed, the Kullback-Leibler divergence between the two models can be written as:

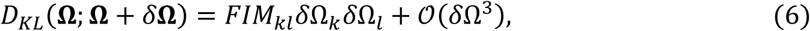

where 1 ≤ *k, l* ≤ 55, and the matrix *FIM* is given by:

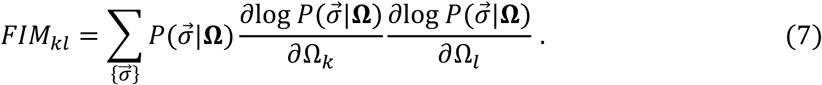

The FIM represents the curvature of the log-likelihood of the model, 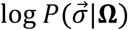, with respect to the model parameters. It quantifies the sensitivity of the model to changes in parameters. By calculating the eigenvalues of the FIM, we can determine which combinations of parameters affect the most the model’s behavior.

In the case of MEM, the FIM can be easily obtained by using equations 1, 2, and 7. As a result, the FIM is given by the covariance matrix of observables associated to the parameters which can be calculated from the model through Monte Carlo simulations, i.e.:

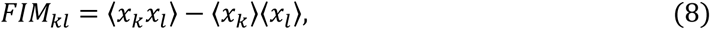

with 1 ≤ *k, l* ≤ 55 and 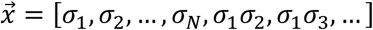.

The FIM was calculated for every neuronal ensemble at every 100-s epoch and it was decomposed into eigenvectors, noted ***v***_*t*,1_, ***v***_*t*,2,_ …, ***v***_*t,k*_, …, where *k* is the rank of the eigenvector and *t* denotes the epoch. We measured the sensitivity of a given parameter by its averaged contribution to the first eigenvector of the FIM, i.e., the sensitivity of *i*-th parameter is given by 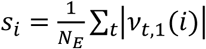, and associated it to the corresponding activity observable (firing rate or correlation). Finally, we separated the activity observables into two classes, called “sloppy” and “stiff”, based on whether the associated sensitivity (*s*) was lower or higher than the median *s*.

### Similarity measures

Temporal variations of model parameters and data statistics were quantified using the average correlation between the parameters/statistics at time t and the parameters/statistics at time *t* + Δ*t*. For example, let 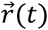 the average firing rates of the neurons during the epoch *t*, the similarity measure is given by:

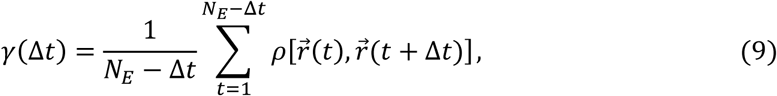

Where *N*_*E*_ is the number of epochs and *ρ* is the Pearson correlation coefficient. In the case of FIM, the matrix was vectorized to calculate *ρ*.

To evaluated how well the data in a given epoch could be explained by the MEM constructed using the data at time *t* + Δ*t*. Specifically, we defined the similarity measure ⟨*D*_*JS*_⟩(Δ*t*), given by the average Jensen-Shannon divergence between the distribution of data binary patterns in epoch *t*, i.e. *P*_data, *t*_, and the distribution of binary patterns predicted by the MEM constructed using the data in epoch *t* + Δ*t*, i.e. *P*_MEM, *t* + Δ*t*_ This measure is given as:

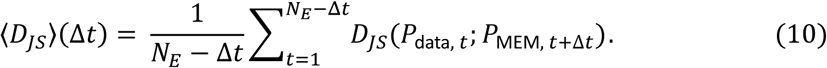

In other words, 1/⟨*D*_*JS*_⟩(Δ*t*) quantifies how well, on average, the model with parameters Ω(*t* + Δ*t*)represents the data from epoch *t*.

### Modulation index

We quantified the responsiveness of the neurons to sensory stimuli through the modulation index (MI) defined as:

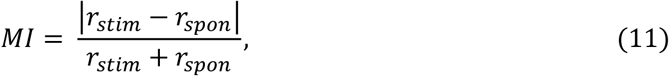

where *r*_*spon*_ is the pre-stimulus average spike count, calculated in the 0.5-s pre-stimulus interval, and *r*_*stim*_ is the average spike count calculated from stimulus onset to 0.5 s after stimulus onset. With this definition, strongly increased or suppressed stimulus responses, with respect to pre-stimulus activity, lead to high MI values.

### Betweenness centrality

For each recording session, we analyzed the network defined by the Pearson correlation matrix of the activities of all single units. The centrality of a neuron, or node, within the network was quantified using the betweenness centrality (BC) measure. BC is given by the number of shortest paths that pass through a given node. The correlation matrix was compute for all 100-s epochs and, for each matrix element, we tested whether the mean of the *N*_*E*_ correlation values differs from 0 (*t* test followed by Bonferroni correction), resulting in a binary graph *G* with entries equal to 1 if correlation were significantly different from zero (corrected p-value < 0.05) and 0 otherwise. The BC for each node of the graph was given by:

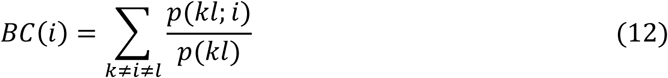

where *p*(*kl*) is the total number of shortest paths from node *k* to node *l* and *p*(*kl*; *i*) is the number of those paths that pass through *i*.

### Neuron-to-population coupling

To quantify the coupling of each neuron to the activity of the neuronal population, we calculated, for each epoch, the Pearson correlation between the activity of each neuron (*σ*_*i*_) and the number of coactive neurons (i.e., with *σ*_*i*_ = +1) at each time bin (*dt* = 10ms) from the neuronal population of single units (without including the neuron *i*). The neuron-to-population coupling was given by the average of the correlation coefficient across epochs.

### ROC analysis

We used the receiver operating characteristic curve (ROC) to evaluate the separation between the distributions of observables from sloppy and stiff classes. Let *X*_sloppy_ and *X*_stiff_ be the sloppy variables, i.e., those variables with associated sensitivity (*s*) lower than the median, and the stiff variables, i.e., those variables with associated sensitivity (*s*) higher than the median *s*, respectively. The ROC curve, *f*(*c*), is build by plotting the probability of *P*(*X*_sloppy_ >*c*) against the probability of *P*(*X*_stiff_ >*c*), for each all *c*. The area under the ROC curve (AUC) is a measure of separation between *P*(*X*_sloppy_)and *P*(*X*_stiff_), and it is given by:

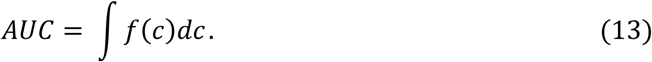

AUC ranges between 0 and 1, with AUC = 0 if *P*(*X*_sloppy_) and *P*(*X*_stiff_) are completely separated and *X*_sloppy_ > *X*_stiff,_ AUC = 1 if *P*(*X*_sloppy_) and *P*(*X*_stiff_) are completely separated and *X*_stiff_ > *X*_sloppy,_, and AUC = 0.5 if *P*(*X*_sloppy_) and *P*(*X*_stiff_) are undistinguishable. We used a permutation test (1000 re-samples), in which observables and classes were randomly associated, to assess AUC values that were significantly different from 0.5.

## Acknowledgements

APA and GD received funding from the FLAG-ERA JTC (PCI2018-092891). GD acknowledges funding from the European Union’s Horizon 2020 FET Flagship Human Brain Project under Grant Agreement 785907 HBP SGA2, the Spanish Ministry Project PSI2016-75688-P (AEI/FEDER) and the Catalan Research Group Support 2017 SGR 1545. GM was supported by a Juan de la Cierva fellowship (IJCI-2014-21937) from the Spanish Ministry of Economy and Competitiveness. AHM received support from the Spanish Ministry of Economy and Competitiveness (BES-2011-049131). JR received funding from the Spanish Ministry of Economy and Competitiveness together with the European Regional Development Fund Grants SAF2010-15730 and SAF2013-46717-R. We thank L. Hollender for sharing her data.

## SUPPLEMENTARY INFORMATION

### Stationary surrogates

To construct the stationary surrogates we first randomly selected a reference epoch *t*. Second, we generated binary data using the MEM estimated from the spiking data at this reference epoch, i.e., using parameters Ω(*t*), through Monte Carlo simulations of the model to obtain 5000 binary patterns. Third, we repeated the Monte Carlo simulations *N*_*E*_ times. Finally, for each of the *N*_*E*_ pieces of surrogate data, we estimated new MEM parameters, Ω′, using gradient descend, and we calculated the corresponding Fisher Information Matrix (FIM). By construction, the obtained surrogate data were stationary and had the same length of the original spiking data. Thus, parameter fluctuations in the surrogate data were only due to estimation errors. Using stationary surrogate data, we found that parameter fluctuations observed in the spiking data were not fully explained by estimation errors and, furthermore, that parameter fluctuations along sloppy dimensions were those that deviated the most from the stationary case (**Figure S1**).

**Figure S1.**
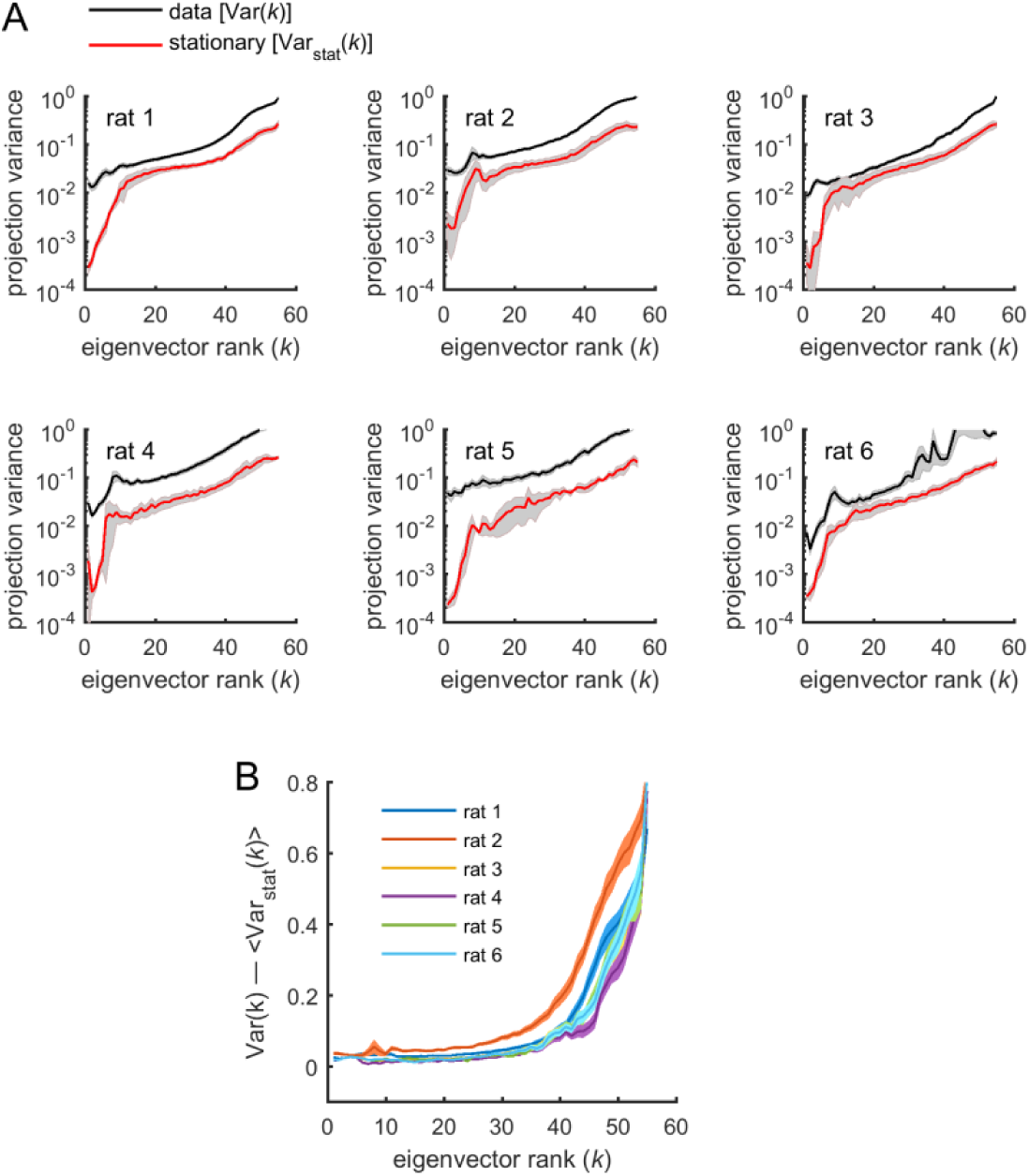
Parameters projections into FIM eigenvectors: data vs. stationary surrogates. Related to Figure 4. The variance of projections of Ω (*t*′) into the different eigenvectors of the Fisher information matrix (FIM) for spiking data was compared to the one obtained from stationary surrogates. **A:** The average variance of projections of model parameters into the different eigenvectors of the FIMs was calculated for the spiking data (parameters Ω(*t*′)) and for the stationary surrogates (parameters Ω′(*t*′)), as a function of the rank *k* of the eigenvectors. Variances were noted Var (*k*) and Var_stat_(*k*) for the spiking data and the stationary surrogates, respectively. Shaded areas represent SEM. Note logarithmic scale on the y-axis. We found that parameter fluctuations were larger than expected by estimation errors in the stationary case for all eigenvectors (i.e., Var(*k*) > Var_stat_(*k*), for all *k*). **B:** The difference between variances of parameter projections estimated from the spiking data and those estimated from stationary surrogates increased with *k*. Thus, parameter fluctuations along sloppy dimensions were those that deviated the most from the stationary case. Shaded areas represent SEM.

### Non-homogeneous Poisson process surrogate data

In Figure 7A-D we found that stiff neurons were more active, more coupled among them, and more central than sloppy neurons. We here investigated the possibility that correlations and BC values could be trivially predicted by firing rates. We tested whether the structure of correlation could arise from spontaneous global fluctuations increasing or decreasing the firing of the neurons, instead of functional connectivity among them. For this, we constructed surrogate data that preserved the mean firing rate of the neurons, and the number and duration of silent (i.e., no spikes, empty 20-ms time bins) and active (i.e., non-empty 20-ms time bins) periods in each epoch, but without other structured correlations among the units.

To do this, we generated non-homogeneous Poisson process (NHPP) surrogate data as follows. We first calculated the average spontaneous firing rates of the single neurons during active periods in the original data, 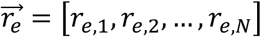, given by the number of spikes, in a given epoch *e*, divided by the total duration of active periods within this epoch. During silent periods the firing rate of all single units was zero. Next, for each unit *i* and within each epoch *e*, we generated non-homogeneous Poisson spike trains using the estimated firing rate as the intensity *λ*_*e,i*_(*t*) of the point process, i.e., *λ*_*e,i*_(*t*) was a step function with *λ*_*e,i*_(*t*) = *r*_*e,i*_, for times *t* within active periods, and *λ*_*e,i*_(*t*) = 0, otherwise. The resulting surrogate data preserved the firing rate of each neuron and silence periods of population activity (**Figure S2A,E**).

Despite being independent during active periods, the units in the NHPP data were correlated due to common global fluctuations across epochs and silent and active periods. Indeed, the mean correlation values in each epoch, ⟨*c*_*ij*_⟩_*e*_, from the original and the surrogate data were highly correlated (**Figure S2B,E**), although ⟨*c*_*ij*_⟩_*e*_ was systematically lower in the surrogate data than in the original data. We tested how well this simple model predicted the structure of correlations, by calculating the pairwise correlations, *c*_*ij*_, and the BC values in the surrogate data, and comparing them to the values obtained in the original data (**Figure S2C-D**). We found that neither *c*_*ij*_ nor BC were predicted by the NHPP: the average fraction of explained variance was equal to R^2^ = 0.148 ± 0.07 and R^2^ = 0.184 ± 0.07 for *c*_*ij*_ and BC, respectively (**Figure S2E**). Moreover, the correlation between firing rates and BC values was substantially reduced in the NHPP, with correlation coefficients equal to 0.59 ± 0.11 and 0.19 ± 0.15 for the original and the NHPP, respectively (p < 0.01, *t*-test). We concluded that correlations and centrality measures were not fully predicted by globally modulated firing rates but were rather suggestive of functional interactions.

**Figure S2.**
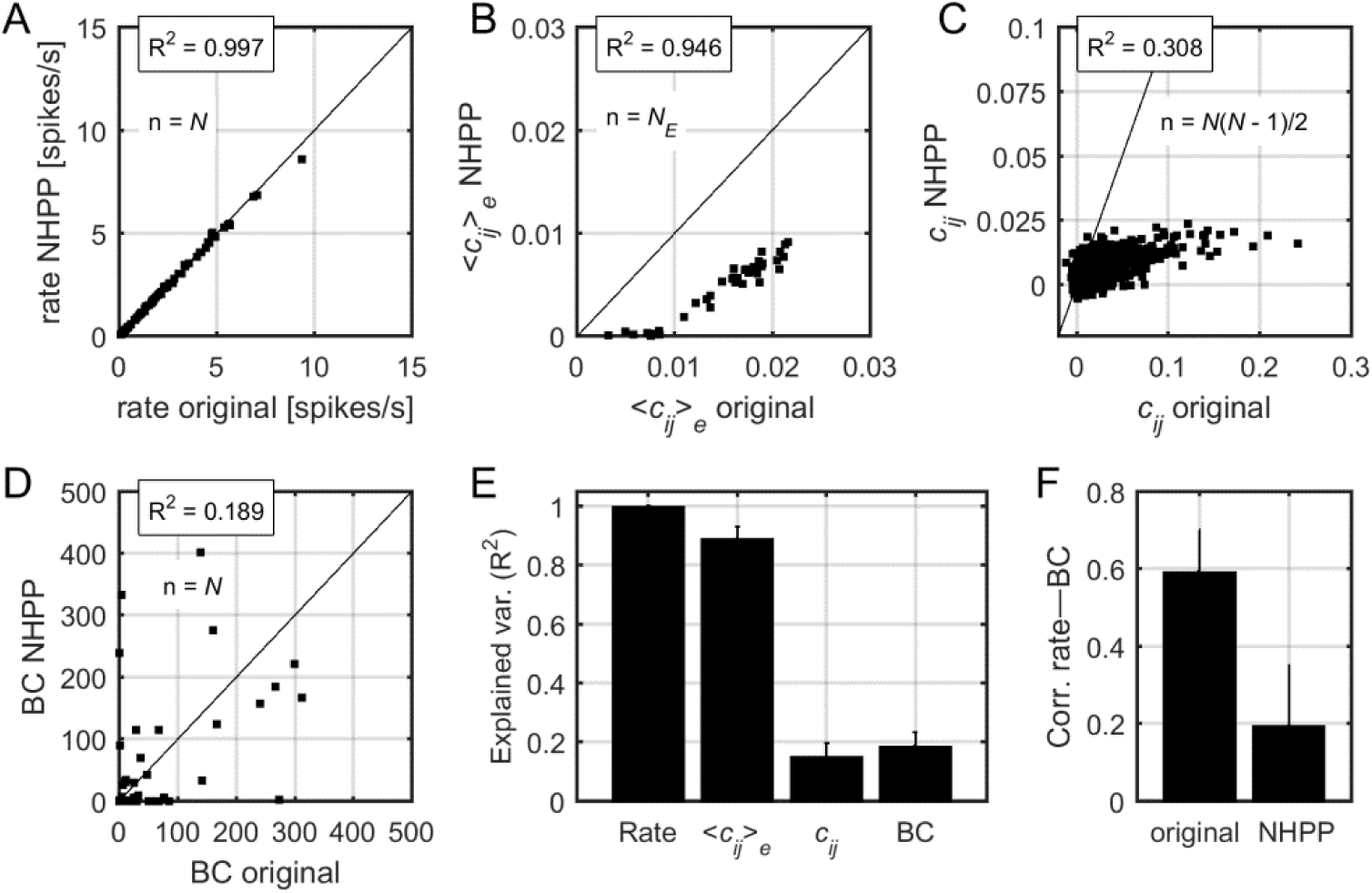
Correlations and centrality in non-homogeneous Poisson process surrogate data. Related to Figure 7. **A:** Relation between firing rates in the original data and firing rates in the non-homogeneous Poisson process (NHPP) surrogates. R^2^ indicates the fraction of explained variance. The black line indicates the identity line. **B:** Relation between the mean pairwise correlation in the original data and that in NHPP surrogates. **C:** Relation between pairwise correlations, averaged over the *N*_*E*_ epochs, in the original data and those in NHPP surrogates. **D:** Relation between BC values in the original data and those in NHPP surrogates (see Methods for calculation of BC). Data in (A), (B), and (C) correspond to one example rat. **E:** Explained variance for firing rates, correlations, and BC values, averaged over datasets. Error bars indicate SEM. **F:** Correlation between firing rates and BC values, averaged over datasets. Error bars indicate SEM.

## References

Arieli A, Sterkin A, Grinvald A, Aertsen A (1996) Dynamics of ongoing activity: Explanation of the large variability in evoked cortical responses. Science 273, 1868–1871. DOI: 10.1126/science.273.5283.1868

Bassett DS, Wymbs NF, Rombach MP, Porter MA, Mucha PJ, Grafton ST (2013) Task-based core-periphery organization of human brain dynamics. PLoS Comput Biol 9, e1003171. DOI: 10.1371/journal.pcbi.1003171

Bazhenov M, Timofeev I, Steriade M, Sejnowski TJ (2002) Model of thalamocortical slow-wave sleep oscillations and transitions to activated States. J Neurosci 22, 8691–8704. DOI: 10.1523/JNEUROSCI.22-19-08691.2002

Beaman CB, Eagleman SL, Dragoi V (2017) Sensory coding accuracy and perceptual performance are improved during the desynchronized cortical state. Nat Commun 8, 1308. DOI: 10.1038/s41467-017-01030-4

Chestek CA, Batista AP, Santhanam G, Yu BM, Afshar A, Cunningham JP, Gilja V, Ryu SI, Churchland MM, Shenoy KV (2007) Single-neuron stability during repeated reaching in macaque premotor cortex. J Neurosci 27, 10742–10750. DOI: 10.1523/JNEUROSCI.0959-07.2007

Compte A, Sanchez-Vives MV, McCormick DA, Wang X-J (2003) Cellular and network mechanisms of slow oscillatory activity (<1 Hz) and wave propagations in a cortical network model. J Neurophysiol 89, 2707–2725. DOI: 10.1152/jn.00845.2002

Csermely P, London A, Wu LY, Uzzi B (2013) Structure and dynamics of core-periphery networks. J Complex Netw 1, 93–123. DOI: 10.1093/comnet/cnt016

Curto C, Sakata S, Marguet S, Itskov V, Harris KD (2009) A simple model of cortical dynamics explains variability and state dependence of sensory responses in urethane-anesthetized auditory cortex. J Neurosci 29, 10600–10612. DOI: 10.1523/JNEUROSCI.2053-09.2009

Destexhe A (2009) Self-sustained asynchronous irregular states and Up-Down states in thalamic, cortical and thalamocortical networks of nonlinear integrate-and-fire neurons. J Comput Neurosci 27, 493–506. DOI: 10.1007/s10827-009-0164-4

Engel TA, Steinmetz NA, Gieselmann MA, Thiele A, Moore T, Boahen K (2016) Selective modulation of cortical state during spatial attention. Science 354, 1140–1144. DOI: 10.1126/science.aag1420

Fruchterman T, Reingold E (1991) Graph Drawing by Force-directed Placement. Software — Practice & Experience. Vol. 21 (11), pp. 1129–1164. DOI: 10.1002/spe.4380211102

Goard M, Dan Y (2009) Basal forebrain activation enhances cortical coding of natural scenes. Nature neuroscience 12(11): 1444–1449. DOI: 10.1038/nn.2402

Hahn G, Ponce-Alvarez A, Monier C, Benvenuti G, Kumar A, Chavane F, Deco G, Frégnac Y (2017) Spontaneous cortical activity is transiently poised close to criticality. PLoS Comput Biol 13, e1005543. DOI: 10.1371/journal.pcbi.1005543

Harris K, Thiele A (2011) Cortical state and attention. Nat Rev Neurosci 12, 509–23. DOI: 10.1038/nrn3084

Hill S, Tononi G (2005) Modeling sleep and wakefulness in the thalamocortical system. J Neurophysiol 93, 1671–1698. DOI: 10.1152/jn.00915.2004

Jercog D, Roxin A, Barthó P, Luczak A, Compte A, de la Rocha J (2017) UP-DOWN cortical dynamics reflect state transitions in a bistable network. eLife 6, e22425. DOI: 10.7554/eLife.22425.001

Kenet T, Bibitchkov D, Tsodyks M, Grinvald A, Arieli A (2003) Spontaneously emerging cortical representations of visual attributes. Nature 425, 954–956. DOI: 10.1038/nature02078

Kitano H (2004) Biological robustness. Nat Rev Genet 5, 826–837.

Luczak A, Barthó P, Harris KD (2009) Spontaneous events outline the realm of possible sensory responses in neocortical populations. Neuron, 62, 413–425.

Luczak A, Barthó P, Harris KD (2013) Gating of sensory input by spontaneous cortical activity. J Neurosci 33, 1684–1695. DOI: 10.1038/nrg1471

Luczak A, Barthó P, Marguet SL, Buzsáki G, Harris KD (2007) Sequential structure of neocortical spontaneous activity in vivo. Proc Natl Acad Sci USA 104, 347–352. DOI: 10.1073/pnas.0605643104

Machta BB, Chachra R, Transtrum MK, Sethna JP (2013) Parameter space compression underlies emergent theories and predictive models. Science 342, 604–607. DOI: 10.1126/science.1238723

MacLean JN, Watson BO, Aaron GB, Yuste R (2005) Internal dynamics determine the cortical response to thalamic stimulation. Neuron 48, 811–823. DOI: 10.1016/j.neuron.2005.09.035

Margolis DJ, Lütcke H, Schulz K, Haiss F, Weber B, Kügler S, Hasan MT, Helmchen F (2012) Reorganization of cortical population activity imaged throughout long-term sensory deprivation. Nat Neurosci 15, 1539–1546. DOI: 10.1038/nn.3240

Marguet SL, Harris KD (2011) State-dependent representation of amplitude-modulated noise stimuli in rat auditory cortex. J Neurosci 31, 6414–6420. DOI: 10.1523/JNEUROSCI.5773-10.2011

Marre O, El Boustani S, Frégnac Y, Destexhe A (2009) Prediction of spatiotemporal patterns of neural activity from pairwise correlations. Phys Rev Lett 102, 138101. DOI: 10.1103/PhysRevLett.102.138101

Mattia M, Sanchez-Vives MV (2012) Exploring the spectrum of dynamical regimes and timescales in spontaneous cortical activity. Cogn Neurodyn 6, 239–250. DOI: 10.1007/s11571-011-9179-4

Mejias JF, Kappen HJ, Torres JJ (2010) Irregular dynamics in up and down cortical states. PLoS One 5, e13651. DOI: 10.1371/journal.pone.0013651

Mochol G, Hermoso-Mendizabal A, Sakata S, Harris KD, de la Rocha J (2015) Stochastic transitions into silence cause noise correlations in cortical circuits. Proc Natl Acad Sci USA 112, 3529–3534. DOI: 10.1073/pnas.1410509112

Okun M, Steinmetz NA, Cossell L, Iacaruso MF, Ko H, Barthó P, et al. (2015) Diverse coupling of neurons to populations in sensory cortex. Nature 521, 511–515. DOI: 10.1038/nature142731

Pachitariu M, Lyamzin DR, Sahani M, Lesica NA (2015) State-dependent population coding in primary auditory cortex. J Neurosci 35, 2058–2073. DOI: 10.1523/JNEUROSCI.3318-14.2015

Panas D, Amin H, Maccione A, Muthmann O, van Rossum M, Berdondini L, Hennig MH (2015) Sloppiness in spontaneously active neuronal networks. J Neurosci 35, 8480–8492. DOI: 10.1523/JNEUROSCI.4421-14.2015

Petersen CCH, Hahn TTG, Mehta M, Grinvald A, Sakmann B (2003) Interaction of sensory responses with spontaneous depolarization in layer 2/3 barrel cortex. Proc Natl Acad Sci USA 100, 13638–13643. DOI: 10.1073/pnas.2235811100

Poulet J, Petersen C (2008) Internal brain state regulates membrane potential synchrony in barrel cortex of behaving mice. Nature 454, 881–5. DOI: 10.1038/nature07150

Sachidhanandam S, Sreenivasan V, Kyriakatos A, Kremer Y, Petersen CCH (2013) Membrane potential correlates of sensory perception in mouse barrel cortex. Nat Neurosci 16, 1671–1677. DOI: 10.1038/nn.3532

Schneidman E, Berry MJ, Segev R, Bialek W (2006) Weak pairwise correlations imply strongly correlated network states in a neural population. Nature 440, 1007–12. DOI: 10.1038/nature04701

Shlens J, Field GD, Gauthier JL, Greschner M, Sher A, Litke AM, Chichilnisky EJ (2009) The structure of large-scale synchronized firing in primate retina. J Neurosci 29, 5022–5031. DOI: 10.1523/JNEUROSCI.5187-08.2009

Tan AYY, Chen Y, Scholl B, Seidemann E, Priebe NJ (2014) Sensory stimulation shifts visual cortex from synchronous to asynchronous states. Nature 509, 226–229. DOI: 10.1038/nature13159

Tkačik G, Marre O, Amodei D, Schneidman E, Bialek W, Berry MJ II (2014) Searching for collective behavior in a large network of sensory neurons. PLoS Comput Biol 10, e1003408. DOI: 10.1371/journal.pcbi.1003408

Tkačik G, Mora T, Marre O, Amodei D, Palmer SE, Berry MJ, et al. (2015) Thermodynamics and signatures of criticality in a network of neurons. Proc Natl Acad Sci 112, 11508–11513. DOI: 10.1073/pnas.1514188112

Transtrum MK, Machta BB, Brown KS, Daniels BC, Myers CR, Sethna JP (2015) Perspective: sloppiness and emergent theories in physics, biology, and beyond. J Chem Phys 143, 010901. DOI: 10.1063/1.4923066

Yassin L, Benedetti BL, Jouhanneau JS, Wen JA, Poulet JF, Barth AL (2011) An embedded subnetwork of highly active neurons in the neocortex. Neuron 68, 1043–50. DOI: 10.1016/j.neuron.2010.11.029

Zagha E, Casale A, Sachdev R, McGinley M, McCormick D (2013) Motor cortex feedback influences sensory processing by modulating network state. Neuron 79, 567–78. DOI: 10.1016/j.neuron.2013.06.008

Zagha E, McCormick D (2014) Neural control of brain state. Curr Opin Neurobiol 10, 178–186. DOI: 10.1016/j.conb.2014.09.010

Ziv Y, Burns LD, Cocker ED, Hamel EO, Ghosh KK, Kitch LJ, El Gamal A, Schnitzer MJ (2013) Long-term dynamics of ca1 hippocampal place codes. Nat Neurosci 16, 264–6. DOI: 10.1038/nn.3329

